# Fronto-parietal Organization for Response Times in Inhibition of Return: The FORTIOR model

**DOI:** 10.1101/134163

**Authors:** Tal Seidel Malkinson, Paolo Bartolomeo

**Affiliations:** INSERM U 1127, CNRS UMR 7225, Sorbonne Universités, and Université Pierre et Marie Curie-Paris 6, UMR S 1127, Institut du Cerveau et de la Moelle épinière (ICM), F-75013 Paris, France

**Keywords:** Inhibition of Return, Attention, Fronto-parietal networks, Right Hemisphere

## Abstract

Inhibition of Return (IOR) refers to a slowing of response times (RTs) for visual stimuli repeated at the same spatial location, as compared to stimuli occurring at novel locations. The functional mechanisms and the neural bases of this phenomenon remain debated. Here we present FORTIOR, a model of the cortical control of visual IOR in the human brain. The model is based on known facts about the anatomical and functional organization of fronto-parietal attention networks, and accounts for a broad range of behavioral findings in healthy participants and brain-damaged patients. FORTIOR does that by combining four principles of asymmetry:

a. Asymmetry in the networks topography, whereby the temporoparietal junction (TPJ) and ventrolateral prefrontal cortex (vlPFC) nodes are lateralized to the right hemisphere, causing higher activation levels in the right intraparietal sulcus (IPS) and frontal eye field (FEF) nodes.
b. Asymmetry in inter-hemispheric connectivity, in which inter-hemispheric connections from left hemisphere IPS to right hemisphere IPS and from left hemisphere FEF to right hemisphere FEF are weaker than in the opposite direction.
c. Asymmetry of visual inputs, stipulating that the FEF receives direct visual input coming from the ipsilateral visual cortex, while the right TPJ and vlPFC and IPS nodes receive input from both the contralateral and the ipsilateral visual fields.
d. Asymmetry in the response modality, with a higher response threshold for the manual response system than that required to trigger a saccadic response. This asymmetry results in saccadic IOR being more robust to interference than manual IOR.

FORTIOR accounts for spatial asymmetries in the occurrence of IOR after brain damage and after non-invasive transcranial magnetic stimulation on parietal and frontal regions. It also provides a framework to understand dissociations between manual and saccadic IOR, and makes testable predictions for future experiments to assess its validity.

**Research highlights:**

- FORTIOR is a model of cortical control of visual IOR in the human brain
- FORTIOR is based on the architecture of fronto-parietal networks
- FORTIOR presents asymmetries favoring the right hemisphere
- FORTIOR explains complex patterns of IOR-related results
- FORTIOR provides testable predictions

## 1 Introduction

Inhibition of Return (IOR) refers to a slowing of response times (RTs) for visual stimuli repeated at the same spatial location, as compared to stimuli occurring at novel locations (Berlucchi, Di Stefano, Marzi, Morelli, & Tassinari, 1981; J. Lupiáñez, Klein, & Bartolomeo, 2006; Posner & Cohen, 1984). In fact, repeated peripheral events can result in faster RTs (facilitation) or slower RTs (IOR), depending on several variables, including the temporal interval between the stimuli, the motor effector used (manual responses or saccades), and the type of visual task (detection or discrimination) (Juan Lupiáñez, 2010). As noted by some theorists (Berlucchi, 2006; Juan Lupiáñez, 2010), this evidence challenges the eponymous account of IOR as inhibition of attention from returning to a previously explored spatial region (Posner, Rafal, Choate, & Vaughan, 1985).

The neural bases of these effects have been the object of extensive research in the last decades. Several lines of evidence indicated an important contribution of the midbrain superior colliculus (SC) in the generation of IOR. For example, in a rare patient with unilateral damage to the right-sided SC, manual IOR was absent only in the visual fields projecting to the damaged SC, i.e., the left temporal hemifield and the nasal right hemifield (Sapir, Soroker, Berger, & Henik, 1999). Consistent with this evidence, recordings in single neurons in the superficial and intermediate layers of the monkey SC showed attenuated activity during IOR (Dorris, Klein, Everling, & Munoz, 2002). However, when saccades were artificially induced by SC microstimulation, IOR reverted to facilitation, with faster saccades to previously stimulated locations. This evidence led Dorris et al. (2002) to conclude that the SC cannot be the site where inhibition is generated; the SC must receive an inhibitory signal from elsewhere, perhaps from the posterior parietal cortex (PPC).

Consistent with this hypothesis, neural activity in monkey LIP was found to be reduced for already explored targets in visual search (Mirpour, Arcizet, Ong, & Bisley, 2009). Also in agreement with the hypothesis of PPC contribution to IOR, in human patients with right hemisphere damage and visual neglect manual IOR for right-sided, non-neglected repeated stimuli was blunted (Bartolomeo, Chokron, & Siéroff, 1999; Bartolomeo, Siéroff, Decaix, & Chokron, 2001), and could even revert to facilitation (Bourgeois, Chica, Migliaccio, Thiebaut de Schotten, & Bartolomeo, 2012). Defective manual IOR was also shown in patients with parietal damage and no signs of neglect (Vivas, Humphreys, & Fuentes, 2003, 2006). An advanced lesion analysis in the Bourgeois et al.’s (2012) study showed that all the patients with reversed manual IOR had damage either to the supramarginal gyrus in the right parietal lobe, or to its connections with the ipsilateral prefrontal cortex. Note, however, that the patients explored by Bourgeois et al. (2012) had normal saccadic IOR.

In addition to these networks, interhemispheric connections can also play a role in the generation of IOR. Case reports on split-brain patients found facilitation instead of IOR when a cued object moved to the contralateral hemifield (Tipper et al., 1997), or slowed appearance of IOR for right-sided targets when the left hemisphere controlled the performance (Berlucchi, Aglioti, & Tassinari, 1997).

Subsequent experiments using Transcranial Magnetic Stimulation (TMS) on normal human participants have provided further evidence concerning the cortical control of IOR. However, the resulting pattern of findings was complex and difficult to reconcile with the simple construct of inhibition of attention to return to a previously stimulated region. This state of affairs provided the motivation for building the present model, which advances a relatively parsimonious proposal restricted to the cortical control of visual IOR in detection tasks. Although the model is primarily based on the TMS evidence reviewed in the following paragraph, it also made use of intracerebral electrophysiological data from human patients and non-human primates.

## 2 TMS interference on IOR

Bourgeois et al. (2013a, 2013b) used repetitive TMS to assess the causal role of distinct nodes of the human fronto-parietal attention networks in the two hemispheres. Participants performed a target-target paradigm (see Maylor & Hockey, 1985). Four black peripheral circles were displayed, at the vertexes of an imaginary square centered on fixation. Participants had to respond to one of the circles becoming white, either by pressing a key or (in a different condition) by making a saccade towards the target. The stimulated brain nodes were the intraparietal sulcus (IPS) and the temporo-parietal junction (TPJ) in each hemisphere. The ensuing complex pattern of results (Table 1) revealed that the TMS effect depended not only on the stimulated node, but also on the presentation side of the visual stimulus (left or right hemifield), and on the response effector. Specifically, TMS on the right hemisphere TPJ decreased IOR only for manual responses with ipsilateral (right) targets, consistent with the patient data (Bourgeois et al., 2012). TMS on the right hemisphere IPS decreased IOR for contralateral (left) targets with both manual and oculomotor responses, but for ipsilateral (right) targets only manual IOR was affected (Bourgeois et al., 2013a). Left hemisphere stimulation had no effect whatsoever on IOR, independent of the stimulated site or of the response effector (Bourgeois et al., 2013b). A further TMS study (Chica, Bartolomeo, & Valero-Cabre, 2011) obtained a similar trend of results in a cue-target paradigm with manual responses, by using double-pulse TMS between cue and target.

**Table 1.** Effects of repetitive TMS stimulation effects on manual and oculomotor IOR (Bourgeois et al., 2013a, 2013b). √, unaffected IOR; X, decreased IOR.

|  | Left-sided targets |  |  |  | Right-sided targets |  |  |  |
| --- | --- | --- | --- | --- | --- | --- | --- | --- |
|  | Left IPS | Left TPJ | Right IPS | Right TPJ | Left IPS | Left TPJ | Right IPS | Right TPJ |
| **Manual response** | √ | √ | X | √ | √ | √ | X | X |
| **Saccadic response** | √ | √ | X | √ | √ | √ | √ | √ |

## 3 The FORTIOR model

This complex pattern of results has no straightforward explanation. To approach this problem, we constructed a model, including the main nodes of the frontoparietal attention networks and the connections between them. The model refers mainly to IOR in target-target detection paradigms, but is relevant for cue-target paradigms as well (see section 3.3.2), and its validity for discrimination tasks is discussed as well (section 5.2). The organizing principles of the model were derived from applying the known anatomical and functional properties of fronto-parietal cortical networks to the behavioral evidence, within the logical constraints necessary for explaining the complex TMS results. The proposed roles of the nodes of the networks are based on evidence from TMS experiments and electrophysiology in humans and non-human primates.

### 3.1 Topography of the model

#### 3.1.1 The FORTIOR nodes

Functional MRI evidence (Corbetta & Shulman, 2002) and tractography results (Thiebaut de Schotten et al., 2011) indicate the existence of fronto-parietal attentional networks, with similar architectures in the monkey and in the human brain (Schmahmann & Pandya, 2006; Thiebaut de Schotten et al., 2011), but with inter-hemispheric asymmetries specific to the human brain (Patel et al., 2015). Schematically, a dorsal attentional network includes the IPS and the FEF, connected by the dorsal branch of the Superior Longitudinal Fasciculus (SLF I). A second, ventral attention network comprises the TPJ/Inferior parietal lobule (IPL) and the ventrolateral prefrontal region (vlPFC; inferior and middle frontal gyri) and is connected by the ventral branch of the SLF (SLF III). Importantly, the SLF I network is thought to be bilateral and symmetric, whereas the SLF III network is strongly lateralized to the right hemisphere (Thiebaut de Schotten et al., 2011). An intermediate branch of the SLF (SLF II) connects the dorsal and ventral attention networks, by traveling from the IPL to the FEF. Based on this architecture, the FORTIOR model includes 6 attention-related nodes: TPJ, IPS, FEF and vlPFC in the right hemisphere; IPS and FEF in the left hemisphere; as well as the right and left visual cortices as visual input entry points (Fig. 1).

**Figure 1.**
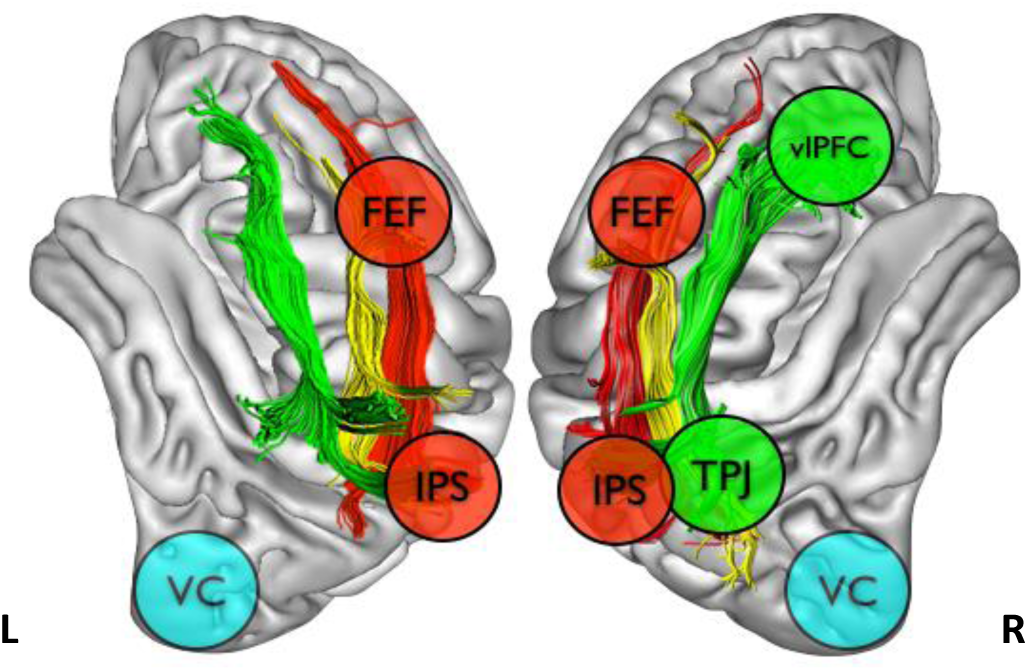
The nodes of the FORTIOR model in the left and in the right hemisphere. FEF, frontal eye field; vlPFC, ventro-lateral prefrontal cortex; IPS, intra-parietal sulcus; TPJ, temporo-parietal junction; VC, visual cortex. Figure modified from Bartolomeo et al (2012).

The model has also two response output nodes: the left motor system for right hand manual responses and the saccade system in both hemispheres for contralateral saccade execution. Note that the response output nodes do not correspond to a single region, directly connected to the other model nodes; rather, they represent response networks.

The nodes have specific roles in the context of the model:

a. **TPJ-** The TPJ is considered here as a hub node, which connects the visual system and other unimodal cortical areas to other attentional nodes (Downar, Crawley, Mikulis, & Davis, 2000; Mars et al., 2012; Wu, Sun, Wang, Wang, & Wang, 2016), as well as ventral and dorsal attentional nodes (Thiebaut de Schotten et al., 2011). It may also play a role in the computations needed for the detection of spatially recurrent events, and in delaying the response to them, thus resulting in behavioral IOR (Ayabe et al., 2008).
b. **vlPFC –** The right vlPFC is important for the detection of relevant targets (Corbetta & Shulman, 2002) and for the generation of a response towards them (Arbula et al., 2017), in the context of an effortful maintenance and execution of a planned behavior (Hampshire, Chamberlain, Monti, Duncan, & Owen, 2010). Perhaps as a consequence of this role in cognitive control, the right vlPFC has also a prominent role in inhibitory processes, such as the generation of stop signals (Aron, Fletcher, Bullmore, Sahakian, & Robbins, 2003; Swann, Tandon, Pieters, & Aron, 2012). In the context of FORTIOR, the main role of the right vlPFC is to generate a signal labeling a stimulus as a target that is task relevant and therefore requires a response.
c. **FEF –** Based on extensive data obtained in humans and in non-human primates, FEF has a double role in the model. First, it controls saccadic responses (Amiez & Petrides, 2009; Paus, 1996). Second, it encodes a priority map of the visual environment (Thompson & Bichot, 2005). In the FORTIOR model, the FEF priority map represents the input to the attention system, rather than the result of the system’s computations, which is encoded in the priority map in the IPS (see section 3.1.1d below). Many studies have shown the involvement of the FEF in spatial attention (Corbetta & Shulman, 2002; Grosbras & Paus, 2002; Thompson & Bichot, 2005), and have associated FEF activity with the processing of stimulus saliency (Thompson & Bichot, 2005), and specifically with IOR (Bichot & Schall, 2002; Mirpour & Bisley, 2015). For example, Bichot and Schall (2002) trained monkeys to perform a saccadic visual search task and found that repetition of target position increased saccade latency and increased the neuronal latency of discrimination between target and distractors. Interestingly, in a similar search task in monkeys, Mirpour and Bisley (2015) identified four different types of neurons in FEF:

- Neurons that responded preferentially to targets over distractors.
- Neurons that responded preferentially when a stimulus that had been fixated was in the response field, thus responding more to repeated targets.
- Neurons that initially showed an enhanced response to a stimulus that had been fixated, but then reversed their response preference 100-150 ms after the saccade.
- Neurons that did not differentiate between search objects, but preferentially responded to the goal of the next saccade. In their data, the first observable response in the FEF upon the fixation of a target was an increased firing rate for repeated versus novel targets. Therefore, we suggest that the FEF map represents at first the occurrence of a salient previous event in the same location as the repeated target. The neurons reversing their preference from previously fixated targets (i.e. repeated targets) to new ones might reflect an activity loop between IPS and FEF, leading to IOR. Mirpour and Bisley (2015) suggest that these data show that reciprocal FEF-IPS processing creates the priority map that guides saccadic eye movements during active, goal-directed visual search. The involvement of FEF in IOR generation was shown not only in visual search tasks, which require some discrimination, but also in detection tasks. For example, using single-pulse TMS over the right FEF during the delay between a peripheral cue and target, Ro et al. (2003) found diminished IOR in the right hemifield, ipsilateral to the TMS. However, there was no measurable IOR modulation when the TMS pulse was applied to the right superior parietal lobule, at variance with the results observed by Bourgeois et al. (2013a) with repetitive TMS on the right IPS.
d. **IPS –** The IPS is considered here as a crucial processing step leading to the delayed response to a target, when the target appears repeatedly at the same spatial location. We suggest that the IPS serves as a priority map encoding the location of the repeated target and the read-out of the saliency signal, which is fed forward to the motor response networks and backward to the visual system. Consequently, the visual system should show reduced activity for repeated targets. Evidence supporting this claim comes from monkey studies showing an increased power in alpha and lower-beta frequencies of local field potentials, together with a decrease in single-neuron responses to previously fixated targets, reflecting an active top-down suppression (Mirpour & Bisley, 2012). Importantly, Miropour et al. (2013) also showed that IPS responses to novel and previously fixated targets start to differ only 60ms after the target-fixation onset. Therefore, IPS is probably not the source of the signal that delays the response to previously fixated targets. Saliency computation might thus be achieved by interactions of the IPS with the FEF, which receives the signals issued from the vlPFC through the TPJ hub. We suggest that these signals are accumulated in the IPS as increased noise in the neuronal population representing the repeated location (see section 3.3.2 below for details). Noise is a ubiquitous property of neural processing, which can be introduced at all stages of the sensorimotor loop and has direct behavioral consequences, from setting perceptual thresholds to affecting movement precision (Faisal, Selen, & Wolpert, 2008). A recent model of bottom-up attention (Khorsand, Moore, & Soltani, 2015) proposed that the formation of saliency signals relies heavily on slow NMDA-mediated recurrent inputs, which simultaneously propagate through successive layers of the network via fast AMPA currents. Computation at successive layers with slow synapses reduces noise and enhances signals such that higher visual areas carry the saliency signals earlier than the lower visual areas. Consequently, feedback from the higher visual areas via fast AMPA synapses can enhance the saliency signals in the lower visual areas. Conversely, enhancing the same noise will result in a noisier saliency signal that will be forwarded to the manual and saccadic motor systems, and delay their response to a repeated target. Here, we suggest that the noise is enhanced in a location-specific manner in the IPS priority map, through an interactive processing between IPS and FEF regions. We further suggest that the sensitivity of the reading response system to the noise is not universal, with the saccadic system being less affected by it than the manual response system (see section 3.2.4 below). Additionally, for manual responses, the IPS read-out is done by the pathway encoding hand movements and requires a transformation from retinotopic to hand coordinates, probably performed in the PPC (Khan, Pisella, & Blohm, 2013). For saccadic responses, the read-out is done via the FEF, through a neuronal population, which is a part of the saccadic control pathway, and does not require coordinate change. This is based on evidence showing that the PPC is engaged in pointing, grasping and reaching, and is involved in spatial representation in reference to hand position (Andersen & Buneo, 2002). According to Anderson and Bueno, the posterior parietal cortex is involved in high-level cognitive functions related to action. These functions include early-movement planning, and particularly the coordinate transformations required for sensory-guided movement. This region is suggested to contain multiple intentional maps with a subdivision to dedicated maps for saccade planning, and different limb movements. These maps are encoded in eye-centered coordinates and include gain-fields underlying the transformation from eye to limb-centered coordinates (also important for the generation of IOR in other coordinate frames, see section 5.4). Thus, attention effects in this region would be related to planning movement. Others see the role of portions of the PPC such as the LIP as closer to attention than to action. For example, LIP activity may be greater before a NOGO response to the saccadic goal than before a GO response (Bisley & Goldberg, 2003). This possibility, however, seems consistent with PPC read-out encoding reduced saliency for specific target locations.

#### 3.1.2 The FORTIOR connections

The connections between the nodes are based on the known connectivity in the human brain:

a. The connections between the attentional nodes are based on the known structure of the SLF in the monkey brain (Schmahmann & Pandya, 2006) and in the human brain (Thiebaut de Schotten et al., 2011). As already mentioned, in the human brain IPS and FEF are connected by a dorsal branch (SLF I), the TPJ and vlPFC are connected by a ventral branch (SLF III) and FEF and TPJ are connected by an intermediate branch (SLF II). There is a gradient of anatomical left-right asymmetries in the human brain. SLF III is generally larger in the right hemisphere than in the left hemisphere, SLF I is symmetrical and SLF II presents a variable degree of right>left asymmetry across individuals (Thiebaut de Schotten et al., 2011). In the present model, for the sake of simplicity, only the right hemisphere SLF III network is considered, given its prominent anatomical (Thiebaut de Schotten et al., 2011) and functional (Corbetta & Shulman, 2002) right>left asymmetry.
b. Local prefrontal connections are the substrate of information transfer between the FEF and vlPFC (Kaufer, 2007; Wood & Grafman, 2003).
c. The connections between early visual cortices and FEF are assumed on the basis of the existence of ultra-fast visual activation in the FEF (Kirchner, Barbeau, Thorpe, Régis, & Liégeois-Chauvel, 2009).
d. In addition, there are bilateral interhemispheric connections between left and right FEF (Catani & Thiebaut de Schotten, 2012; Kaufer, 2007) and between the left and right IPS (Catani & Thiebaut de Schotten, 2012; Koch et al., 2011). As mentioned before, case reports on split-brain patients (Berlucchi, Aglioti, & Tassinari, 1997; Tipper et al., 1997) suggest a role for inter-hemispheric connections in IOR.
e. Callosal connections between the ventral nodes (TPJ and vlPFC) are instead less prominent (Catani & Thiebaut de Schotten, 2012, see their Fig. 9.4): these nodes work in relative isolation from their contralateral homologues.

### 3.2 Organizing principles

The model is organized around four principles. Some of these principles are supported by existing evidence, while others are more speculative.

#### 3.2.1 Asymmetrical network topography

The dorsal SLF I network (connecting the nodes IPS and FEF) is relatively symmetrical across the hemispheres. The ventral SLF III network (TPJ and vlPFC) is instead lateralized to the right hemisphere (Corbetta & Shulman, 2002; Thiebaut de Schotten et al., 2011). The right-lateralization of the SLF III network induces a certain degree of functional asymmetry between the right and left SLF I networks, because only the right hemisphere SLF I network receives direct additional stimulation from the ventral SLF III network (Gigliotta, Malkinson, Miglino, & Bartolomeo, 2017).

#### 3.2.2 Inter-hemispheric connectional asymmetry

There is an asymmetry in the inter-hemispheric white fibers connecting the dorsal fronto-parietal nodes, such that information transmission through inter-hemispheric connections from the left hemisphere to the right hemisphere is weaker and slower than in the opposite direction (Marzi, 2010). There is TMS-based evidence that this is indeed the case for inter-parietal connections (Koch et al., 2011). A similar bias was put forth in an fMRI-based model of attention networks in which there was an asymmetry in the strength of connections between bilateral IPS with preference of the right-to-left connection (Siman-Tov et al., 2007). Additionally, electrophysiological and behavioral studies suggest that a relative abundance of fast-conducting myelinated axons in the right hemisphere might be the cause of both a right hemispheric activation increase and a faster signal transfer from the right to the left hemisphere (Barnett & Corballis, 2005). For the present purposes, we shall assume that some connectional asymmetry of this type also exists between the FEFs.

#### 3.2.3 Asymmetrical visual inputs

The model stipulates that the FEF receives direct visual input coming from the ipsilateral visual cortex, while the right TPJ and vlPFC receive input from both the contralateral and the ipsilateral visual fields. The IPS nodes are activated both for contralateral and ipsilateral targets, through intra-hemispheric and inter-hemispheric connections. Preliminary evidence obtained from intracerebral recordings in patients with drug-resistant epilepsy confirms that the right IPS can respond to targets presented in both visual fields (Seidel Malkinson et al., 2017). Specifically, electrodes recording from the right IPS showed an IOR-related validity effect not only for contralateral targets but also for ipsilateral ones.

#### 3.2.4 Response modality asymmetry

Motor output relies on partially distinct network dynamics, depending on the used effector. IPS activity influences manual responses through M1 and premotor cortex (Filimon, 2010), and saccadic responses though the FEF (Buschman & Miller, 2007). Moreover, we put forth that the saccade network is more encapsulated (i.e., less prone to interference), and has a lower threshold for response initiation, than the manual response system. Thus, saccade initiation is faster and more automatic than manual responses. This feature is in line with studies reporting a dissociation between manual and saccadic response patterns within the same task (Bompas, Hedge, & Sumner, 2017). Also, saccadic responses can be immune to visual illusions which influence manual responses (Lisi & Cavanagh, 2015, 2017). Lisi and Cavanagh (2017) accounted for this difference by suggesting that the saccade system relies on a representation that accumulates visual information and location errors over shorter time windows than the representation used for controlling hand movements (see further details in section 3.3.2). In the present model, the shorter integration window is implemented as a lower signal to noise (SNR) threshold of the saccadic response (i.e. greater tolerance to noise), which causes its earlier and more reliable production, compared to manual responses. As described above (section 3.1.1d), we suggest that the delayed response in IOR results from increased noise in the IPS priority map, which the response networks read out and act upon. As a result, the response networks require additional time to process the noisier representation of a repeated target in the IPS priority map and to trigger a response. In this context, a lower SNR threshold for the saccadic response means that a shorter computation time and a smaller priority signal are needed for the saccade system to read out the map and trigger a response for a repeated target. If so, then saccadic IOR should appear at an earlier stimulus onset asynchrony (SOA) than manual IOR. Indeed, saccadic IOR typically occur at an SOA of around 100-200ms (Klein, 2000), while manual IOR tends to appear at a later SOA of 200-300ms (Samuel & Kat, 2003). Further support for the higher efficiency of the saccadic IOR response relies on the relatively more direct pathway leading from the FEF to saccade execution regions than the one connecting the IPS to motor regions controlling manual responses. The FEF is linked directly, and indirectly through the superior colliculus, to the paramedial pontine reticular formation, and from there to the oculomotor cranial nerve nuclei that control saccades (Kandel, Schwartz, Jessell, Siegelbaum, & Hudspeth, 2000). On the other hand, the IPS connections are more indirect, passing through two intricate parieto-frontal networks: the inferior and the superior networks. The inferior network connects the IPS to many other parietal regions such as the anterior intraparietal region and to the ventral premotor area and to the motor cortex; the superior network, which might be lateralized to the left in humans (Stark & Zohary, 2008), connects it to a hub in the medial wall of PPC, with strong connections to premotor regions and to the motor cortex (Grafton, 2010; Karl & Whishaw, 2013). This difference between IPS and FEF in their connectivity to motor command regions contributes to the difference in the weight of the priority map in the response output: activation in the IPS map needs to be stronger and longer to delay a manual response, than to delay the saccadic response through the FEF.

As a results of these constraints, the model supposes an asymmetry between left and right hemisphere IPS and FEF, whereby the left hemispheric nodes drive a weaker output that is insufficient to delay the manual response and trigger a manual IOR by itself, but remains sufficient for the generation of a saccadic IOR due to the saccadic system’s lower threshold and greater noise tolerance.

### 3.3 The temporal sequence of information flow in the model

This section describes the temporal sequence of information flow in the FORTIOR model, when a visual target appears repeatedly at the same location and entails an IOR response, as described in Bourgeois et al.’s TMS experiments (Bourgeois et al., 2013a, 2013b), as well as in cue-target paradigms (Chica, Martin-Arevalo, Botta, & Lupianez, 2014). Because TMS effect on IOR changed according to the stimulus presentation side, we modeled separately left-sided and right-sided visual presentation.

#### 3.3.1 Registering the occurrence of a first target

In order to delay the response toward a repeated target, the system must first know that a target is repeated. Thus, some kind of trace must be kept of the first target identity and location that will modulate the response toward a subsequent target, appearing at the same location. FORTIOR suggests that this trace is kept in the FEF priority map, which reflects the saliency of the environment, perhaps in the form of a baseline shift in the firing rate of the neuronal population representing the stimulated location in the visual field (Figure 2). Specifically, the visual activation triggered by the appearance of the first target is transferred from the visual cortex (VC) to the FEF in the same hemisphere. Because TMS interference in a cue-target paradigm on the right hemisphere TPJ decreased IOR after left-sided cues, but not after right-sided cues (Chica et al., 2011), we suggest that visual information arrives at the right FEF also via the right TPJ for left-sided targets, but not for right-sided targets. Visual input generates a location-specific activation in the priority maps in the FEF and exchanged with the IPS (Figure 2). Evidence for baseline increases during directed endogenous attention to a cued location and in the absence of visual stimulation have been found using monkey electrophysiology (Colby, Duhamel, & Goldberg, 1996; Luck, Chelazzi, Hillyard, & Desimone, 1997) and in human fMRI studies (Kastner, Pinsk, De Weerd, Desimone, & Ungerleider, 1999). Importantly, this increased baseline-firing rate was observed *before* the appearance of the visual stimulus. In an electrophysiology study in monkeys, attention increased consistently the baseline-firing rate of V4 and V2 cells, about 175ms after the monkeys were cued by a predictive peripheral cue to a location inside the neurons’ receptive fields compared to a location outside the receptive fields (Luck et al., 1997). There was little or no effect on the peak stimulus-evoked response. In human fMRI studies, it was shown that baseline increases occurred across the visual system (Kastner et al., 1999). The FEF was suggested to be a possible source of these baseline shifts, as it was found to have greater baseline increases than ventral stream areas and the IPS, and to reflect the attentional demands of the task rather than the sensory processing (Kastner et al., 1999). Based on this evidence, here we propose that the FEF might signal the occurrence of a salient exogenous event using a similar mechanism.

**Figure 2.**
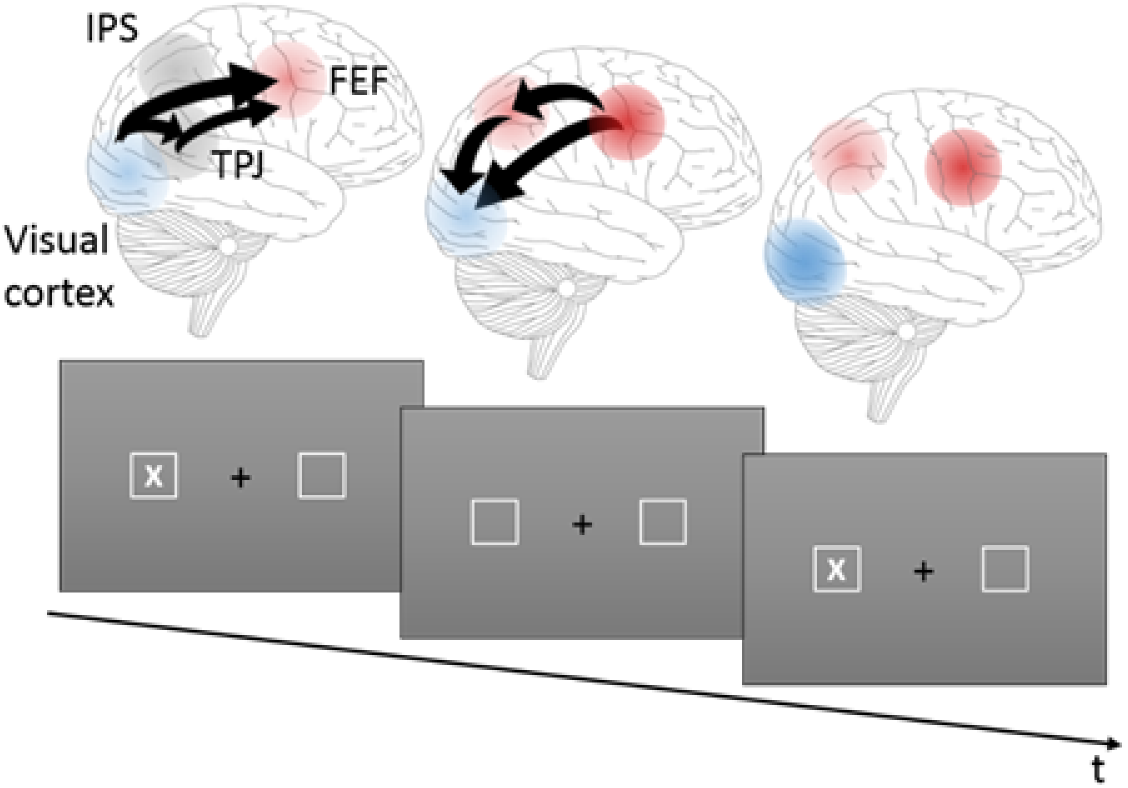
A schematic illustration of the registration of the occurence of the first target (X in bottom panel). Left Visual activation (depicted in blue) related to the first target is tranferred from the visual cortex to the FEE directly for left-sided or right-sided targets, or also through TPJ for left-sided targets, causing saliency-related FEF activation (depicted in red). Middle: The baseline activation in the neural population representing the spatial location in the FEF priority map shifts (depicted as a deeper red). The baseline shift is transferred to the IPS map (depicted in red) and coveyed to the visual cortex (in deeper blue). Right: Baseline shifts persist in the delay period until the onset of the second target.

#### 3.3.2 Responding to repeated stimuli

The IOR-inducing stimulus can be a repeated target in target-target paradigms, or a cued target in cue-target paradigms. In cue-target paradigms, a cue toward which no response should be made is first presented. This lack of response to the cue allows a speeded presentation of cue and target, impossible otherwise. The cue is detected and its location registered in the FEF priority map (as described in the previous section 3.3.1); any inappropriate response to the cue must be inhibited, perhaps by right prefrontal regions (Aron, 2004 #4674, but see Criaud & Boulinguez, 2013; Erika-Florence, Leech, & Hampshire, 2014). With short SOAs (less than 200-300ms for manual responses, and less than 100-200ms for saccadic responses), the activation in the FEF priority map caused by the cue may be summed up with the activation caused by the subsequent target, creating a stronger signal with an early onset time (see Lupiáñez, 2010). This may reflect the limited temporal resolution of the response systems: Within a time window of 100-200ms only a single saccade can be performed, whereas in 200-300ms only a single manual response can be made. Therefore, there could be only a single response toward multiple stimuli processed during these temporal windows. Consequently, a hard-wired constraint of the system might be to treat such compressed set of stimuli as if they belonged to a single event, and to sum them up in the priority map. Such summation would increase the activation related to the target and thus lead to facilitation, with faster RT for validly cued targets. Possibly, this facilitation might reflect an error in detection, whereby the target is not detected as a separate event from the cue, an error that can lead to effects like the Illusory Line Motion (Chica, Charras, & Lupiáñez, 2008), or accelerated Temporal Order Judgements (Schettino, Rossi, Pourtois, & Müller, 2016). However, with longer SOAs, when two discrete responses can be made toward successive stimuli, the stimuli will not be considered as a single event but as two separate ones, and their related activations in the priority maps will not be summed up. Instead, it is advantageous to maximize the separation between them, to ensure an appropriate response toward each stimulus.

We suggest that outside this summation window, the previously activated location in the priority maps will accumulate noise. Noise can be defined as trial-to-trial variability (when the stimulus is held constant) and can result from two sources. The first source is the specific properties of the system. For example, the initial state of the neural circuitry may be different at the beginning of each trial, resulting in different neuronal and behavioral responses (Faisal et al., 2008). Baseline changes may be at the basis of such a change in the initial state of the network. The variability in the response will increase if the system's dynamics are highly sensitive to the initial conditions (Faisal et al., 2008). The second source of variability is random or irregular fluctuations or disturbances that are not part of a signal, or simply put - noise. This noise is generated at all stages of the visual neural processing, from the photoreceptor level, until the synapse and network levels. The two sources are not independent. For example, the generation of an action potential is highly sensitive to noise when the membrane potential is near the firing threshold (Faisal et al., 2008). Thus, if the baseline shifts up and noise is added, the trial-to-trial variability will increase, and the representation of information will be less reliable. In general, the less the information represented, the more the evidence needed to reach a perceptual decision and the longer the processing time required.

One way noise could affect the IPS-FEF maps is by increasing baseline firing rate (without increasing the responses to stimuli), which reduces the signal to noise ratio (SNR) at the specific location in the IPS priority map and therefore represents a lower saliency signal for a given stimulus strength.

Adding baseline noise to the priority map in the previously activated location will filter out weak, unreliable signals appearing at the same place (which might be residual decaying activation of the previous stimulus), and promote the processing of only strong, salient stimuli that appear there. Thus, in a target-target paradigm, the SNR of the activation in the priority map related to the target repeated at the same location will be smaller compared to that caused by the first target. This weaker activation will be propagated to the regions driven by the output of the map. Indeed, previous studies in monkeys have found a reduction of visual responses at previously cued locations in SC neurons (Dorris, Klein, Everling, & Munoz, 2002; Robinson & Kertzman, 1995), PPC (Constantinidis & Steinmetz, 2001; Robinson, Bowman, & Kertzman, 1995; Steinmetz, Connor, Constantinidis, & McLaughlin, 1994), LIP (Mirpour et al., 2009) the inferior temporal cortex (Miller, Gochin, & Gross, 1991), and PFC (DeSouza & Everling, 2004).

Additionally, at the network level, noise may affect the representation of information in maps, like those in FEF and IPS, by altering the correlation of noise between neurons. Maps represent information in a distributed code and have information capacity that may be greatest when the noise sources across the population are not correlated (Averbeck, Latham, & Pouget, 2006; Zohary, Shadlen, & Newsome, 1994). Thus, increases in noise correlation, for example, can potentially affect the encoding of information in those maps, and make their readout more demanding and requiring more processing time. Relatedly, Cohen and Maunsell (Cohen & Maunsell, 2009) studied how spatial attention affected noise correlation by recording simultaneously from dozens of neurons in V4 region in monkeys, while monkeys performed an orientation change detection task in which the target was preceded by a valid spatial cue. Results showed that a reduction in noise correlation between the neurons accounted for the majority of the attentional improvement in population sensitivity, more than the increase in the neurons’ firing rate. Thus, attentional effects can be explained by noise modulation at the network level. Yet, as far as we know, the involvement of noise modulation in the generation of IOR has not been directly tested, and remains speculative for now.

The next sections describe FORTIOR’s account of the processing of the repeated target that leads to a delayed response.

##### 3.3.2.1 The flow of visual information related to the repeated target

Visual input regarding the recurring stimulus originates from visual cortices in striate regions (labeled VC in Fig 3a) contralateral to the stimulus, and is transferred to the ipsilateral FEF and to the right hemisphere TPJ. Information about right-sided visual stimuli, causing left visual cortex activation, spreads from the left FEF to its right counterpart. However, the left-to-right inter-hemispheric link is relatively weak (assumption 2.2.2.), and therefore contributes to a relatively small feed-forward wave into the right vlPFC.

**Figure 3.**
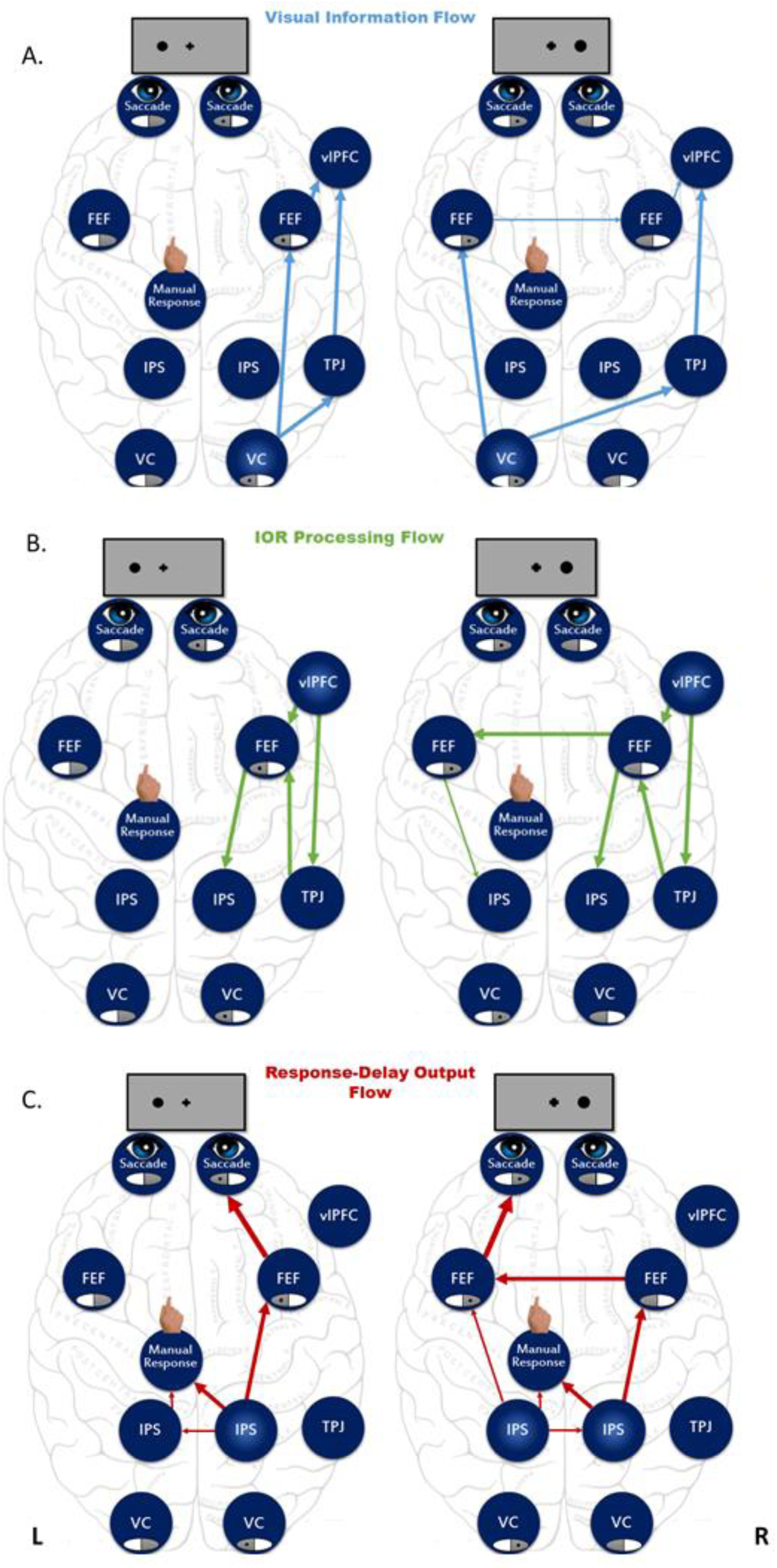
The temporal sequence of information flow in FORTIOR. A. Visual information (blue arrows) about the repeating left sided (left panel) and right sided stimulus (right panel) is conveyed from the contralateral visual cortices to the right TPJ and contralateral FEF, and then to the right vIPFC. B. The right vIPFC initiates the processing of the IOR and sends forward through a network comprising the TPJ and FEF, a delaying signal that enhances the noise in the IPS priority map through IPS-FEF interaction (green arrows; left sided stimuli - left panel; right sided stimuli - right panel). C. The noisy location specific representation in the IPS map is read more slowly by the motor system and causes a delayed manual response and by the FEF for a delayed saccadic response (red arrows; left sided stimuli - left panel; right sided stimuli -right panel). Arrow thickness represents connection strength.

##### 3.3.2.2 IOR processing flow

When visual information generated by the spatially recurrent stimulus arrives at the vlPFC node, vlPFC labels the stimulus as a target requiring a response, and initiates a signal that is sent forward through a network comprising the TPJ and FEF. The FEF priority map already encoding the previously activated location as a baseline shift, now processes the incoming vlPFC input and transfers it to the IPS as location specific increased noise (Fig 3b). Additionally, the relayed vlPFC signal may possibly cause a decrease in the noise correlation in neuronal populations encoding the location of the repeated target in the FEF-IPS maps. For left-sided targets, this IOR-related processing only implicates right hemisphere nodes. When the target is on the right, the signal is also sent through to the left FEF, and from there to the left IPS.

##### 3.3.2.3 Response-delay output flow

Perturbing the nodes in the network in the Bourgeois et al.’s (2013a) study caused a decrease in IOR, i.e. faster RTs, and not a diminished response *per se*; thus, under normal conditions the FORTIOR output, coming from the IPS priority map, must lead to a delayed response. The IPS map output is transferred to the motor system for manual responses, and through the FEF for saccadic responses (Fig 3c). The response networks require more time to read out the noisy representation of the second target in the IPS map, which holds less information, and thus the response is delayed. For left-sided targets, the right IPS co-activates the left IPS and both trigger a delay of the manual response. The left IPS also transfers the map output to the right FEF, which causes a delayed leftward saccade. For right-sided targets, both right and left IPS send the priority map output to the motor system, with a greater weight assigned to the right IPS contribution (see above, section 3.2.4). The IPS map in both hemispheres is also read by the respective interconnected FEF regions, but only the left FEF can initiate a delayed saccadic command for saccades to right-sided targets, because the left hemisphere attention networks have predominantly contralateral competence.

## 4 FORTIOR accounts for TMS effects on IOR

### 4.1 TMS on the right hemisphere IPS

In the Bourgeois et al.’s (2013a) study, TMS on the right IPS caused a reduction in both manual and saccadic IOR for left-sided targets, but only in manual IOR for right-sided targets (see Table 1). The present model accounts for this dissociation by invoking a disruption of the priority map output, encoded by the stimulated right IPS node (Fig. 4). TMS stimulation interferes with the location-specific enhancement of noise in the map; consequently, noise is not increased for repeated targets, which are processed as if they were new ones. With left-sided targets, the right IPS is the only source of the priority map output, and thus, its disruption results in diminished manual and saccadic IOR. With right-sided targets, due to the retinotopic organization of the visual cortex, left hemispheric nodes are also involved, and the left IPS participates in the generation of the priority map output. However, the left IPS output is weak and on its own is insufficient to drive a manual delayed response. This is due to the lack of direct input from the right ventral nodes (because of weak callosal connections, see assumption 3.1.2.e), causing a weaker activation levels in the left IPS. On the other hand, this weak signal is sufficient to delay the saccadic response, initiated via the FEF, due to the lower threshold of the saccadic system.

**Figure 4.**
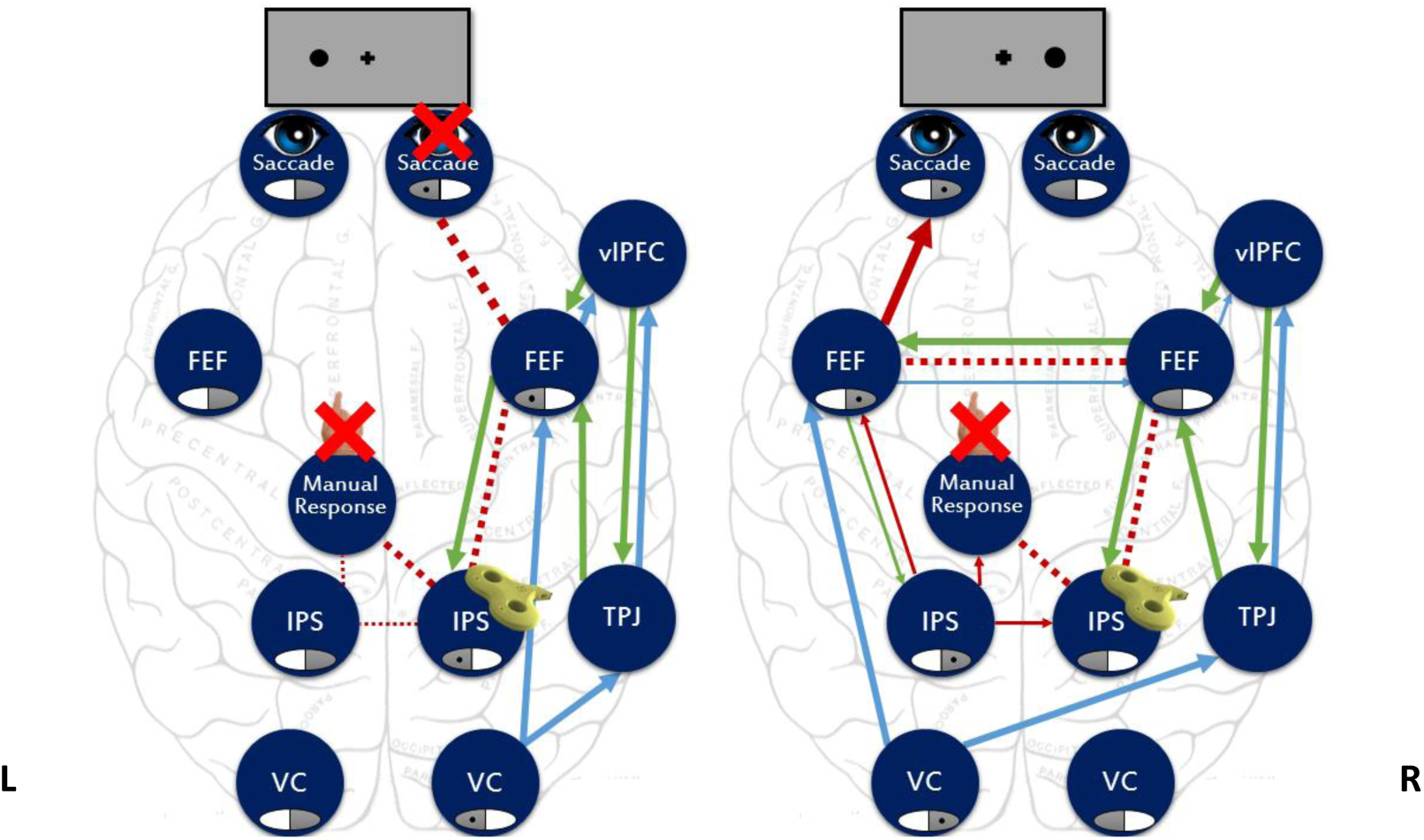
FORTIOR account for the pattern of IOR reduction after TMS stimulation of the right hemisphere IPS. TMS affects the enhancement of location-specific noise in the priority map in the right IPS (red dashed lines), leaving unaffected the visual information (blue arrows) and IOR processing (green arrows) flows. For left-sided targets (left panel) this disruption perturbs both manual and saccadic IOR, because the right IPS is the only source for the priority map noisy output. For right-sided targets (right panel), residual weak noise is encoded in the left IPS map (red full arrows), which is sufficient for delaying the saccadic response, but not the manual response.

### 4.2 TMS on the left hemisphere IPS

Stimulating the left IPS with TMS did not affect either manual or saccadic IOR for right- or left-sided targets (Bourgeois et al., 2013b, see Table 1). Within the present model, the right hemisphere IPS is the main involved node for the attentional processing of left-sided targets. Thus, interference on the left hemisphere IPS has no influence on IOR (Fig. 5). With right-sided targets, the left IPS is also read by the motor response networks, but its priority map generates a relatively weak output. For manual response to be delayed, the right IPS, which responds to both left-sided targets and right-sided targets, needs to back up this weak activation. When the left IPS is disrupted, the read-out from the right IPS is sufficient for manual IOR to occur. Similarly, the noisy representation in the right hemisphere IPS map is sufficient to delay the saccadic response.

**Figure 5.**
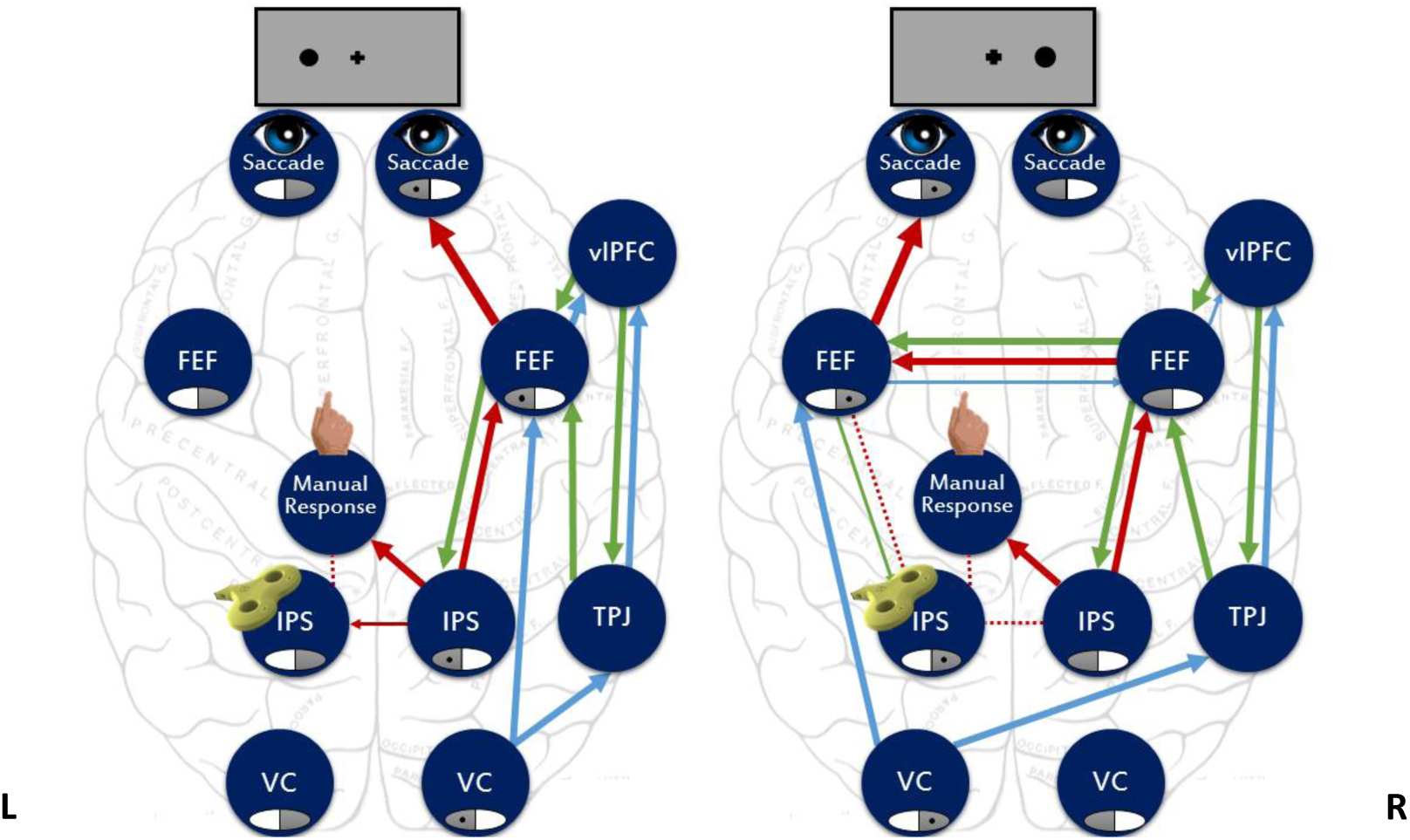
FORTIOR account for the absence of IOR reduction after TMS stimulation of the left hemisphere IPS. TMS affects the generation of noisy location-specific representation in the left IPS priority map (red dashed lines); however, this leaves unaffected the noisy priority map in the right hemisphere IPS (red arrows), the visual information (blue arrows) and the IOR processing (green arrows). For left-sided targets (left panel), the right hemisphere IPS is the main node involved, and thus the disruption of the left hemisphere IPS has no effect. For right-sided targets (right panel), the output of the right hemisphere IPS triggers both a saccadic and manual IOR.

### 4.3 TMS on the right hemisphere TPJ

TMS interference on the right TPJ caused a reduction only in manual IOR for right-sided targets (Bourgeois et al., 2013a, see Table 1). In the framework of FORTIOR, right TPJ stimulation disrupts the transmission of ipsilateral visual input through the TPJ hub to the right vlPFC, and consequently disrupts the signal from the vlPFC to the right FEF. This vlPFC signal labels the repeated stimulus as a task-relevant target requiring response (Fig. 6). The disrupted transmission entails the interruption of noise accumulation in the right IPS via FEF-IPS interactions, and disrupts the prolonged reading of the IPS noisy map by the response networks. However, for left-sided targets, the transfer of contralateral visual information through the right FEF to the vlPFC and back allows to circumvent the disruption, and produce a strong enough activation in the right IPS for generating both manual and saccadic IOR responses. For right-sided targets, the weak inter-hemispheric transfer of ipsilateral visual information through the FEFs to the vlPFC is sufficient only for the generation of a weak signal in the right vlPFC, which fails to trigger the noise enhancement FEF-IPS loop in the right hemisphere. However, the weak vlPFC output travels back to the left FEF-IPS loop, enabling the enhancement of noise in the left IPS map. The weak left IPS noisy map is then read out slower by the saccadic response system, via the left FEF, and delays the saccadic response towards the right-sided target. Nevertheless, the activation in the left IPS priority map is too weak to delay the manual response system, which treats the target as a novel one, with consequent lack of manual IOR.

**Figure 6.**
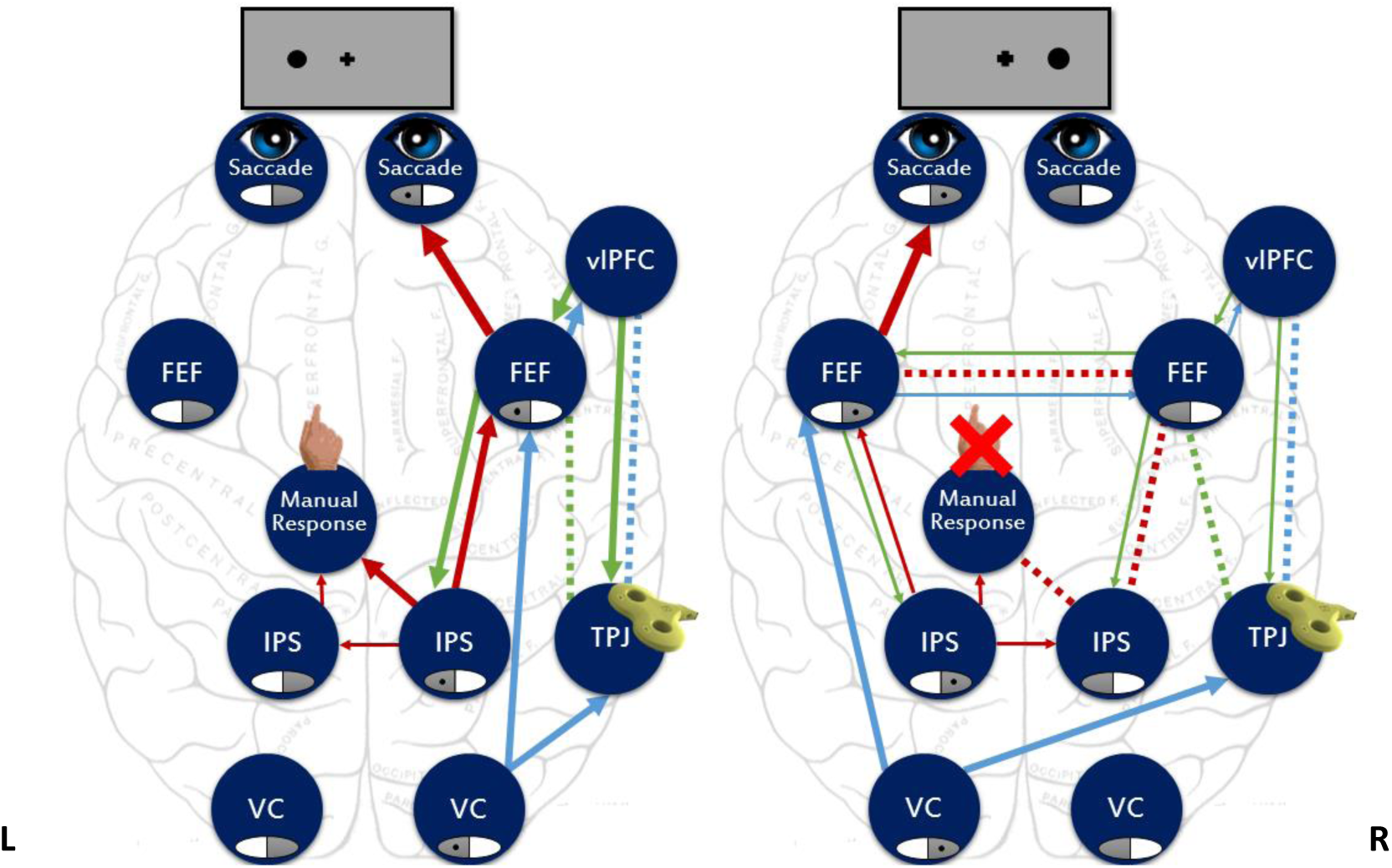
FORTIOR account for the pattern of IOR reduction after TMS stimulation of the right hemisphere TPJ. TMS disrupts the transmission of ipsilateral visual input (blue dashed line) through the TPJ hub to the right vlPFC, and of the signal from the vlPFC to the right hemisphere FEF (dashed green line). The disrupted transmission causes a failure of labeling the repeated stimulus as a relevant target, and entails the interruption of the location-specific accumulation of noise in the right hemisphere IPS (red dashed lines). For left-sided targets (left panel), the transfer of contralateral visual information (blue arrows) through the right hemisphere FEF to the vlPFC and back, allows to circumvent the disruption, and leads to location-specific noise in the IPS map that is sufficient for generating both manual and saccadic responses. For right-sided targets (right panel), the weak rightward inter-hemispheric transfer of ipsilateral visual information through the FEFs to the vlPFC is sufficient only for activating a weak signal (green thin arrow). This weak signal fails to trigger a noise enhancement loop in the right hemisphere FEF-IPS priority maps. As a consequence, the repeated target is treated as a novel one by the manual response system. However, transmission of the same weak vlPFC signal from the right FEF to the left FEF (green thin arrow) is sufficient to trigger a noise enhancement FEF-IPS loop in the left hemisphere. Due to its lower threshold, the saccadic response system can read out the noisy left IPS priority map, and trigger a delayed rightward saccade (normal saccadic IOR).

### 4.4 TMS on the left hemisphere TPJ

TMS on the left TPJ did not interfere with IOR (Bourgeois et al., 2013b). According to the present model, this is explained by the fact that the left TPJ is not involved in the processing of IOR, and is not a part of the relevant network.

## 5 Discussion

The aim of FORTIOR is to provide a theoretical model explaining the cortical basis of IOR generation in detection paradigms that would also account for the complex pattern of interference produced by TMS stimulation on IPS and TPJ (Bourgeois et al., 2013a, 2013b). Thus, FORTIOR completes and extends previous models of IOR based on SC functioning (Satel, Wang, Trappenberg, & Klein, 2011), which explicitly called for further modeling of cortical contributions to IOR.

FORTIOR does that by combining four principles of asymmetry:

e) Asymmetry in the networks topography, whereby the TPJ and vlPFC nodes are lateralized to the right hemisphere, causing higher activation levels in the right IPS and FEF nodes.
f) Asymmetry in inter-hemispheric connectivity, in which inter-hemispheric connections from left IPS to right IPS and from left FEF to right FEF are weaker than in the opposite direction.
g) Asymmetry of visual inputs, stipulating that the FEF receives direct visual input coming from the ipsilateral visual cortex, while the right TPJ and vlPFC and IPS nodes receive input from both the contralateral and the ipsilateral visual fields.
h) Asymmetry in the response modality, with a higher response threshold for the manual response system than that required to trigger a saccadic response. This asymmetry results in saccadic IOR being more reliable and robust to interference than manual IOR.

FORTIOR suggests that IOR results from enhanced noise in the representation of the repeatedly stimulated location in the priority map in IPS. Enhanced noise lowers the SNR when the map is read out by the response systems. The noisy map requires more processing time, which leads to a delayed response. When the enhancement of noise is perturbed, for example by TMS-based interference on the IPS, the repeated target is treated as if it were a novel one. According to FORTIOR, IOR is a result of interactions in a network of nodes; the model therefore predicts that perturbing non-redundant nodes (such as lateralized nodes) and their connections will decrease IOR, following the patterns predicted by the above-mentioned asymmetries in the model.

FORTIOR is based on evidence from human and monkey electrophysiology and human neuroimaging studies, with particular focus on the constraints introduced by the results of two repetitive TMS stimulation studies (Bourgeois et al., 2013a, 2013b). Yet, it is important to keep in mind that such studies rely on a limited number of subjects and experiments, and that there is debate on the duration of TMS effect and on TMS influence on remote interconnected areas (Eisenegger, Treyer, Fehr, & Knoch, 2008). Therefore, FORTIOR remains a suggested framework for the cortical control of IOR that needs to be further assessed and refined with new data in future studies. A number of predictions can be generated based on FORTIOR to test its validity.

### 5.1 Testable model predictions

#### 5.1.1 IOR in Visual Neglect

Visual neglect provides an example for a condition in which right cortical lesions accompany abnormal IOR. Lesions associated with neglect typically affect the caudal nodes of the FORTIOR model in the right hemisphere, of their white matter connections to the frontal nodes (Bartolomeo et al., 2012). Neglect patients often show blunted manual IOR or even facilitation (faster RTs) for repeated right-sided, non-neglected stimuli (Bartolomeo, Chokron, & Siéroff, 1999; Bartolomeo, Siéroff, Decaix, & Chokron, 2001; Bourgeois, Chica, Migliaccio, Thiebaut de Schotten, & Bartolomeo, 2012). However, when saccadic responses were tested, the same patients had normal saccadic IOR for the same right-sided repeated targets (Bourgeois et al., 2012). Moreover, neglect patients tended to show normal manual and saccadic IOR to left-sided targets. Advanced lesion analysis showed that all the patients with reversed manual IOR in the Bourgeois et al.’s study (2012) had damage to the supramarginal gyrus in the right parietal lobe, or to its connections with the ipsilateral prefrontal cortex. Yet, the link between neglect and abnormal IOR might be more complex. Vivas et al. (Vivas et al., 2003, 2006) tested patients with parietal damage even in the absence of neglect signs, and found that they also demonstrated decreased IOR (but not facilitation) on the ipsilesional side. These data stress that manual IOR may strongly depend on cortical networks including the right parietal lobe, which are typically dysfunctional in neglect patients (Bartolomeo, Thiebaut de Schotten, & Doricchi, 2007; Doricchi, Thiebaut de Schotten, Tomaiuolo, & Bartolomeo, 2008; He et al., 2007; Mort et al., 2003; Thiebaut de Schotten et al., 2005). Importantly, a number of patients with right brain damage can eventually compensate for clinical signs of neglect (Lunven et al., 2015), while still showing signs of spatial bias on more stringent tests (Bartolomeo, 1997, 2000; Bonato, 2012). This could have been the case for at least some of the parietal patients tested by Vivas et al. (2003, 2006). Moreover, in the Vivas et al’s studies eye movements were not controlled; if patients directed their gaze at ipsilesional cues or initial targets (a common event in right brain-damaged patients: Bourgeois et al., 2015; Gainotti, D'Erme, & Bartolomeo, 1991), the second stimulus was presented on their fovea; then fast responses to foveal stimuli could have offset IOR. Finally, the level of detail of the anatomical analysis of lesions in these studies was insufficient to draw firm conclusions about the identity of the cortical circuits implicated in the modulation of IOR.

In the frame of FORTIOR, neglect patients with damage to the supramarginal gyrus in the right parietal lobe, or to its connections with the ipsilateral prefrontal cortex (see Bourgeois et al., 2012) have lesions in regions corresponding to the right TPJ node and its connections to the right vlPFC (SLF III network). Thus, in the framework of FORTIOR these lesions lead to a failure to transfer visual information via the TPJ to the vlPFC, similar to TMS stimulation of the right TPJ (see above section 4.3). For right sided targets, due to the asymmetry in visual inputs and in interhemispheric connections, this leads to a failure in triggering noise enhancement in the right hemisphere FEF-IPS loop and to the absence of location specific noise in the right hemisphere IPS priority map. The location specific noisy representation of the repeated target in the left IPS is sufficient to delay the saccadic response for right-sided targets, but is too weak to be read by the manual response system, causing the absence of manual IOR. The paradoxical facilitation of the manual responses to right-sided targets in these patients might reflect an abnormal persistence of the initial enhanced activation in the FEF priority map in response to the first target, causing a summation of the activation of the successive co-localized targets. Some support for the existence of such prolonged integration window in neglect patients comes from the study of attentional blink in those patients, showing that on top of abnormal spatial attention, they also have abnormal temporal attention dynamics (Husain, Shapiro, Martin, & Kennard, 1997). Specifically, when detecting two letter targets, serially embedded in a stream of rapidly presented visual stimuli at a single location, neglect patients with right parietal, frontal or basal ganglia lesions failed to detect the second target when the delay between the targets was three times as long as for healthy controls and patients without neglect. Based on these data, Husain and colleagues (1997) suggested that once attention is committed to the analysis of a visual object, neglect patients’ ability to direct their attention to another object is impaired, even when these are presented at the same location. This impairment can contribute to an abnormal persistence of the activation for the first target, leading to the paradoxical facilitation found in those patients (Bartolomeo et al., 1999, 2001; Bourgeois et al., 2012). FORTIOR predicts that patients with selective right hemisphere IPS damage will show patterns of performance similar to TMS interference on right IPS (see section 4.1 and fig 3 above).

#### 5.1.2 TMS-based disruption of right hemisphere vlPFC will perturb IOR

Because of its role in detecting task-relevant targets and generating responses toward them, and due to its lateralization to the right hemisphere, perturbing the functioning of the right hemisphere vlPFC is predicted to cause a failure to identify the second target as a task-relevant one, and thus to prevent the triggering of the FEF-IPS noise enhancement loop. As a result, the repeated target will be treated by the response systems as a novel one, diminishing both manual and saccadic IOR.

#### 5.1.3 TMS-based disruption of the FEF will perturb IOR

Since according to FORTIOR the FEF is crucial for the both the registering of the occurrence of the first target and for the triggering of the location specific enhancement of noise in the IPS priority map, its disruption by TMS should affect IOR generation. In FORTIOR the visual input to the FEF is suggested to come from the ipsilateral visual cortex and from the contralateral FEF. Furthermore, because the right-to-left inter-hemispheric connection is stronger and because the dorsal nodes in the right hemisphere are suggested to have stronger activation, the disruption is predicted to affect IOR differentially according to the side of the target. FORTIOR predicts that perturbing the right FEF by repetitive TMS will affect both the registration of the occurrence of the first target, and the accumulation of location specific noise in the IPS priority map upon the occurrence of the second target.

#### 5.1.4 FEF is activated between the first and the second target

According to FORTIOR the occurrence of the first target is registered in the FEF, possibly as a baseline activation shift. This has been demonstrated using fMRI (Kastner et al., 1999), but should be tested with more direct measures such as intracerebral recordings.

#### 5.1.5 Noise enhancement in the IPS priority map

The model suggests that noise is enhanced in the IPS specifically at the repeated target location. Thus, measurements of IPS activity should show a decrease in SNR in location-specific neuronal populations and/or in the noise correlation in those neural populations. For example, neural activity in monkey PPC recordings or human intracerebral recordings should show increased trial-to-trial variability for repeated targets.

#### 5.1.6 Callosal connections are essential for IOR

FORTIOR suggests that callosal connections are important for IOR generation, especially for right-sided targets. Split-brain patients provide a potential source of data to test this suggestion. As already mentioned, Tipper et al. (1997) described that two split-brain patients that despite having normal IOR within each visual field, showed facilitation instead of IOR when a cued target moved to the opposite visual field. Tipper et al. suggested that a subcortical cross-hemispheric transfer of facilitation and a callosal transfer of IOR could explain this pattern. This is an object-based IOR, in which cue and target co-occur in the same location relative to the object, unlike the simpler scenario of IOR in the same retinotopic location, as described in FORTIOR. Yet, in the framework of FORTIOR, these data could reflect the need for a right vlPFC contribution to generate IOR, which then needs to be transferred across hemispheres when the cue and the target are on different visual fields. Without the contribution of the vlPFC, there would be no detection of the repeated target as relevant and no enhancement of noise in the representation of the IPS-FEF priority maps. Possibly, under these conditions, the repeated activation at the cued and target object location would be summed together, resulting in a facilitation-like phenomenon. However, in the Tipper et al.’s study, results for right- and left-sided targets were presented together, thus potential hemifield differences are not visible. As a matter of fact, another split brain patient, studied by Berlucchi et al. (1995), had blunted/delayed IOR in his right hemispace, controlled by the left hemisphere, consistent with the dominance of right hemisphere networks for IOR suggested by FORTIOR. Another option for testing this prediction is by using intraoperative stimulation of white matter fibers (see Thiebaut de Schotten et al., 2005). If these connections are indeed important for the generation of IOR, then stimulating them should change IOR in a hemifield-dependent manner.

### 5.2 Cortical control of IOR in discrimination tasks

The FORTIOR model provides an account for cortical control of visual IOR in detection tasks. However, IOR is also evident in discrimination tasks, albeit at longer cue-target intervals (Lupiáñez, Milan, Tornay, Madrid, & Tudela, 1997). Discrimination tasks require more complex visual processing than detection tasks, which might be carried out at different cortical regions depending on the exact nature of the task, and therefore entail more complex cortical interactions than the detection tasks described in FORTIOR. Yet, Lupiáñez and his co-workers (Lupiáñez, Martín-Arévalo, & Chica, 2013) suggested that the same cognitive mechanisms underlying IOR in detection tasks are also responsible for IOR in discrimination tasks. According to the Detection Cost Theory of IOR (Lupiáñez, 2010; Lupiáñez et al., 2013), the IOR effect reflects a cost of rapidly detecting or encoding the appearance of new objects or events when they are similar to previous attention-capturing events. According to this framework, cueing a location hinders the detection of a subsequent target at the very same location, whereas it facilitates selecting this same target for subsequent perceptual discriminative processing leading to its recognition. In contrast to detection tasks, with discrimination the cueing effect is usually less negative or even positive due to another opposite effect: spatial selection benefit. Considering this, FORTIOR might provide a general explanation for the IOR effect (detection cost) that leads to the behavioral observation of IOR. IOR would occur prominently in detection tasks, and only when the other opposite positive effect of cueing is cancelled in discrimination tasks (e.g., Martín-Arévalo, Chica, & Lupiáñez, 2014).

### 5.3 Motor and Perceptual components of IOR

A central debate in IOR literature concerns the existence of motoric and perceptual components in IOR. This distinction can relate to the causes of IOR or to its effects (Lupiáñez, Klein, & Bartolomeo, 2006). Regarding the latter, it seems that different experimental designs and task demands can emphasize one component over the other, and that IOR may operate at several stages of processing to discourage orienting toward previously cued locations (Lupiáñez et al., 2006). For example, to test this dichotomy Taylor and Klein (2000) used central arrows and peripheral events as the first or second of two successive stimuli. Responses to the same direction indicated by the first stimulus were delayed even when the second stimulus was a central arrow when the oculomotor system was involved. In contrast, when the oculomotor system was not engaged and only manual responses were made to the second stimulus (after the first one was ignored, or responded to manually), they found attentional/perceptual IOR. These findings may suggest that the motor/perceptual distinction corresponds, at least partially, to the division between saccadic and manual IOR in FORTIOR (section 3.2.4).

### 5.4 The reference frame of IOR

FORTIOR focuses on retinotopic IOR, as this is the initial and simplest form of spatial representation in the visual system. However, the reference frame of IOR is a contentious question, based on conflicting evidence. In general, an extra-retinal reference frame (spatiotopic: such as a head-based, egocentric or object-based reference frames) is needed to deal with image stability problems (caused by eye movements for example), which are not properly dealt with by the initial retinotopic representation of visual stimuli is. IOR was reported to occur in both retinotopic and spatiotopic coordinates (Krüger & Hunt, 2013), initially in retinotopic and only later in spatiotopic coordinates (Mathôt & Theeuwes, 2010), or in spatiotopic coordinates (Pertzov, Zohary, & Avidan, 2010; Posner & Cohen, 1984; Satel, Wang, Hilchey, & Klein, 2012). This complex pattern of evidence necessitates further investigations and meta-analyses. Several potential (and not mutually exclusive) neural mechanisms were suggested to be at the basis of nonretinotopic visual representation, and all of them could possibly serve nonretinotopic IOR:

1. Gain fields – First described in neurons in the posterior parietal cortex by Andersen and Mountcastle (1983). Retinotopically organized neurons modulate their visual response according to gaze-position. The readout of the neuronal population can generate a spatiotopic representation in the absence of explicit individual neural spatiotopic responses (R. A. Andersen & Zipser, 1988; Zipser & Andersen, 1988, see also 3.1.1d).
2. Predictive Remapping – According to this view, there is no higher order spatial map in the brain but instead the representation of the visual world always remains in retinotopic coordinates (Wurtz, 2008; Wurtz, Joiner, & Berman, 2011). At a neuronal level, the updating of a retinotopic map is achieved by predictive remapping - a mechanism by which the receptive field (RF) of a neuron is shifted towards its future retinotopic post-saccadic position, slightly before the saccade execution. Neurons that show evidence of shifting RFs have been studied in LIP, where they were initially discovered (Colby, Duhamel, & Goldberg, 1996; J. R. Duhamel, Colby, & Goldberg, 1992; Heiser & Colby, 2006), and have subsequently been found in the FEF (Umeno & Goldberg, 1997, 2001). Shifting RFs have also been reported (though in a smaller percentage of the neurons) in earlier extra-striate visual areas (Nakamura & Colby, 2002; Tolias et al., 2001).
3. Remapping of attentional pointers – Here, the updating is done on the post-saccadic position of a few objects of interest in the visual field, and not of the entire visual field. This activation transfer serves as an early warning to cells whose receptive fields are about to receive an attended target, therefore enabling stability across eye movements (Cavanagh, Hunt, Afraz, & Rolfs, 2010). Spatial attention shares spatial maps with saccade control centers in areas like superior colliculi, LIP and FEF and the activity peaks on these maps guide saccades and index the location of the target on other similarly organized retinotopic maps throughout the brain. Activity in these attention/saccade maps can activate the corresponding locations in other regions, enhancing processing for information from the corresponding target location in these areas.
4. Real Position (Spatiotopic) Neurons – Image stability is achieved by explicit spatiotopic neurons that respond to stimuli in one region of visual space rather than one point on the retina. These spatiotopic “real position” neurons were found in the parietal cortex (Galletti, Battaglini, & Fattori, 1993) and in the ventral intraparietal area (J.R. Duhamel, Bremmer, BenHamed, & Graf, 1997). In general, real position neurons have been found in areas that overlap those where the gain field neurons are found (Galletti & Fattori, 2002). However, real position neurons are rather rare.

In the face of conflicting evidence regarding the frames of reference of IOR and the multitude of hypotheses regarding the underlying neural mechanisms, we feel that currently there is not enough data to establish a generalized model explaining the generation of IOR in the different proposed frames of reference. Nevertheless, most of the cortical regions found to be involved in coordinates transformation by the different suggested mechanisms, are also modeled as nodes in FORTIOR, making it amenable to explain the generation of IOR in different frames of reference.

### 5.5 IOR in other modalities

Besides vision, IOR was also reported in touch and audition, and between all cue – target pairings of vision, touch, and audition (Spence, Lloyd, McGlone, Nicholls, & Driver, 2000). Since IOR is a spatial phenomenon, i.e. RT is longer for previously stimulated locations, its exact multi-modal nature may depend on the nature of the spatial information that the different senses convey and their underlying neural mechanisms. Vision, audition and touch vary in their coordinate frame of space (e.g. vision is retinotopic, touch is somatotopic, etc.) and/or their spatial resolution (vision has the best spatial resolution). For example, tactile IOR was shown to be modulated by the somatotopic rather than by the allocentric distance between the cued body site and the target (Röder, Spence, & Rösler, 2002). Thus, IOR in audition and touch probably depends in part on distinct sensory brain regions that encode space in modality-specific frames of reference. Likewise, FORTIOR nodes with retinotopic mapping, such as the FEF, might participate primarily in the generation of visual IOR. Yet, some of the regions involved in visual IOR, such as the TPJ and vlPFC nodes, are probably supra-modal and participate in IOR generation in all modalities as well as across-modalities. Support for this idea comes from two event-related fMRI experiments in which the right TPJ and right inferior frontal gyrus responded to changes in sensory stimuli in visual, auditory and tactile modalities (Downar et al., 2000), and their response was stronger for novel rather than familiar multimodal stimuli (Downar, Crawley, Mikulis, & Davis, 2002). These data suggest that the right TPJ and inferior frontal gyrus are a part of a multimodal network for involuntary attention to events in the sensory environment. Another potential multimodal node, the posterior parietal cortex, was shown to contain multimodal spatial maps (R.A. Andersen, Snyder, Bradley, & Xing, 1997). For example, Macaluso et al. (2000) tested the effect of simultaneous visuo-tactile stimulation on the activity of the human visual cortex. They found that tactile stimulation enhanced activity in the visual cortex when it was on the same side as a visual target, and that this crossmodal spatial attention was mediated via back-projections from multimodal parietal areas. Therefore, FORTIOR’s IPS node may be also involved in the generation of cross-modal IOR.

In conclusion, here we have presented a model of bi-hemispheric, fronto-parietal cortical control of IOR, which takes into account a large amount of evidence from monkey neurophysiology, human neuroimaging and non-invasive brain stimulation, and makes specific predictions allowing to assess its validity in future research.

## Acknowledgements

Supported by ANR BRANDY R16139DD (PB), by fellowships of the Israel Science Foundation 57/15 (TSM) and Marie Skłodowska-Curie 702577 (TSM). The authors wish to thank Juan Lupiáñez and an anonymous reviewer for their very helpful comments on a previous version of this article.

## References

Andersen, R. A., & Mountcastle, V. B. (1983). The influence of the angle of gaze upon the excitability of the light-sensitive neurons of the posterior parietal cortex. Journal of Neuroscience, 3(3), 532–548.

Andersen, R. A., Snyder, L. H., Bradley, D. C., & Xing, J. (1997). Multimodal representation of space in the posterior parietal cortex and its use in planning movements. Annual Review of Neuroscience, 20, 303–330.

Andersen, R. A., & Zipser, D. (1988). The role of the posterior parietal cortex in coordinate transformations for visual–motor integration. Canadian journal of physiology and pharmacology, 66(4), 488–501.

Aron, A. R., Fletcher, P. C., Bullmore, E. T., Sahakian, B. J., & Robbins, T. W. (2003). Stop-signal inhibition disrupted by damage to right inferior frontal gyrus in humans. Nature Neuroscience, 6(2), 115–116.

Averbeck, B. B., Latham, P. E., & Pouget, A. (2006). Neural correlations, population coding and computation. Nature reviews. Neuroscience, 7(5), 358.

Ayabe, T., Ishizu, T., Kojima, S., Urakawa, T., Nishitani, N., Kaneoke, Y., & Kakigi, R. (2008). Neural processes of attentional inhibition of return traced with magnetoencephalography. Neuroscience, 156(3), 769–780.

Bartolomeo, P. (1997). The novelty effect in recovered hemineglect. Cortex, 33(2), 323–332.

Bartolomeo, P. (2000). Inhibitory processes and compensation for spatial bias after right hemisphere damage. Neuropsychological Rehabilitation, 10(5), 511–526.

Bartolomeo, P., Chokron, S., & Siéroff, E. (1999). Facilitation instead of inhibition for repeated right-sided events in left neglect. NeuroReport, 10(16), 3353–3357.

Bartolomeo, P., Siéroff, E., Decaix, C., & Chokron, S. (2001). Modulating the attentional bias in unilateral neglect: The effects of the strategic set. Experimental Brain Research, 137(3/4), 424–431.

Bartolomeo, P., Thiebaut de Schotten, M., & Doricchi, F. (2007). Left unilateral neglect as a disconnection syndrome. Cerebral Cortex, 45(14), 3127–3148.

Bartolomeo, P., De Schotten, M. T., & Chica, A. B. (2012). Brain networks of visuospatial attention and their disruption in visual neglect. Frontiers in human neuroscience, 6.

Berlucchi, G., Aglioti, S., & Tassinari, G. (1997). Rightward attentional bias and left hemisphere dominance in a cue-target light detection task in a callosotomy patient. Neuropsychologia, 35(7), 941–952.

Bonato, M. (2012). Neglect and extinction depend greatly on task demands: a review. Front Hum Neurosci, 6, 195. doi: 10.3389/fnhum.2012.00195

Bourgeois, A., Chica, A. B., Migliaccio, R., Bayle, D. J., Duret, C., Pradat-Diehl P., … Bartolomeo, P. (2015). Inappropriate rightward saccades after right hemisphere damage: Oculomotor analysis and anatomical correlates. Neuropsychologia, 73, 1–11. doi: 10.1016/j.neuropsychologia.2015.04.013

Bourgeois, A., Chica, A. B., Migliaccio, R., Thiebaut de Schotten, M., & Bartolomeo, P. (2012). Cortical control of inhibition of return: Evidence from patients with inferior parietal damage and visual neglect. Neuropsychologia, 50(5), 800–809.

Bourgeois, A., Chica, A. B., Valero-Cabré, A., & Bartolomeo, P. (2013a). Cortical control of inhibition of return: causal evidence for task-dependent modulations by dorsal and ventral parietal regions. Cortex, 49(8), 2229–2238. doi: 10.1016/j.cortex.2012.10.017

Bourgeois, A., Chica, A. B., Valero-Cabré, A., & Bartolomeo, P. (2013b). Cortical control of Inhibition of Return: exploring the causal contributions of the left parietal cortex. Cortex, 49(10), 2927–2934. doi: 10.1016/j.cortex.2013.08.004

Catani, M., & Thiebaut de Schotten, M. (2012). Atlas of the Human Brain Connections: Oxford University Press.

Cavanagh, P., Hunt, A. R., Afraz, A., & Rolfs, M. (2010). Visual stability based on remapping of attention pointers. Trends Cogn Sci, 14(4), 147–153. doi: S1364-6613(10)00028-8 [pii] 10.1016/j.tics.2010.01.007

Chica, A. B., Charras, P., & Lupiáñez, J. (2008). Endogenous attention and illusory line motion depend on task set. Vision Research, 48(21), 2251–2259.

Chica, A. B., Martin-Arevalo E., Botta, F., & Lupianez, J. (2014). The Spatial Orienting paradigm: How to design and interpret spatial attention experiments. Neurosci Biobehav Rev, 40c, 35–51. doi: 10.1016/j.neubiorev.2014.01.002

Cohen, M. R., & Maunsell, J. H. (2009). Attention improves performance primarily by reducing interneuronal correlations. Nature neuroscience, 12(12), 1594–1600.

Colby, C. L., Duhamel, J. R., & Goldberg, M. E. (1996). Visual, presaccadic, and cognitive activation of single neurons in monkey lateral intraparietal area. Journal of Neurophysiology, 76(5), 2841–2852.

Constantinidis, C., & Steinmetz, M. A. (2001). Neuronal responses in area 7a to multiple stimulus displays: II. Responses are suppressed at the cued location. Cerebral Cortex, 11(7), 592–597.

Corbetta, M., & Shulman, G. L. (2002). Control of goal-directed and stimulus-driven attention in the brain. Nature Reviews Neuroscience, 3(3), 201–215.

Criaud, M., & Boulinguez, P. (2013). Have we been asking the right questions when assessing response inhibition in go/no-go tasks with fMRI? A meta-analysis and critical review. Neuroscience & biobehavioral reviews, 37(1), 11–23.

Doricchi, F., Thiebaut de Schotten, M., Tomaiuolo, F., & Bartolomeo, P. (2008). White matter (dis)connections and gray matter (dys)functions in visual neglect: Gaining insights into the brain networks of spatial awareness. Cortex, 44(8), 983–995.

Dorris, M. C., Klein, R. M., Everling, S., & Munoz, D. P. (2002). Contribution of the primate superior colliculus to inhibition of return. Journal of Cognitive Neuroscience, 14(8), 1256–1263.

Downar, J., Crawley, A. P., Mikulis, D. J., & Davis, K. D. (2000). A multimodal cortical network for the detection of changes in the sensory environment. Nature Neuroscience, 3(3), 277–283. doi: 10.1038/72991

Downar, J., Crawley, A. P., Mikulis, D. J., & Davis, K. D. (2002). A cortical network sensitive to stimulus salience in a neutral behavioral context across multiple sensory modalities. Journal of Neurophysiology, 87(1), 615–620.

Duhamel, J. R., Bremmer, F., BenHamed S., & Graf, W. (1997). Spatial invariance of visual receptive fields in parietal cortex neurons. Nature, 389(6653), 845–848.

Duhamel, J. R., Colby, C. L., & Goldberg, M. E. (1992). The updating of the representation of visual space in parietal cortex by intended eye movements. Science, 255(5040), 90–92.

Erika-Florence M., Leech, R., & Hampshire, A. (2014). A functional network perspective on response inhibition and attentional control. Nature communications, 5.

Faisal, A. A., Selen, L. P., & Wolpert, D. M. (2008). Noise in the nervous system. Nature reviews. Neuroscience, 9(4), 292.

Gainotti, G., D’Erme P., & Bartolomeo, P. (1991). Early orientation of attention toward the half space ipsilateral to the lesion in patients with unilateral brain damage. Journal of Neurology, Neurosurgery and Psychiatry, 54, 1082–1089.

Galletti, C., Battaglini, P. P., & Fattori, P. (1993). Parietal neurons encoding spatial locations in craniotopic coordinates. Experimental brain research, 96(2), 221–229.

Galletti, C., & Fattori, P. (2002). Posterior parietal networks encoding visual space. In H. O. Karnath, A. D. Milner & G. Vallar (Eds.), The Cognitive and Neural Bases of Spatial Neglect (pp. 59–62). Oxford, UK: Oxford University Press.

Grosbras, M.-H., & Paus, T. (2002). Transcranial magnetic stimulation of the human frontal eye field: effects on visual perception and attention. Journal of cognitive neuroscience, 14(7), 1109–1120.

He, B. J., Snyder, A. Z., Vincent, J. L., Epstein, A., Shulman, G. L., & Corbetta, M. (2007). Breakdown of functional connectivity in frontoparietal networks underlies behavioral deficits in spatial neglect. Neuron, 53(6), 905–918.

Heiser, L. M., & Colby, C. L. (2006). Spatial updating in area LIP is independent of saccade direction. Journal of Neurophysiology, 95(5), 2751–2767.

Husain, M., Shapiro, K., Martin, J., & Kennard, C. (1997). Abnormal temporal dynamics of visual attention in spatial neglect patients. Nature, 385(6612), 154–156.

Koch, G., Cercignani, M., Bonnì, S., Giacobbe, V., Bucchi, G., Versace, V., … Bozzali, M. (2011). Asymmetry of parietal interhemispheric connections in humans. J Neurosci, 31(24), 8967–8975. doi: 10.1523/jneurosci.6567-10.2011

Krüger, H. M., & Hunt, A. R. (2013). Inhibition of return across eye and object movements: The role of prediction. Journal of Experimental Psychology: Human Perception and Performance, 39(3), 735.

Lunven, M., Thiebaut De Schotten, M., Bourlon, C., Duret, C., Migliaccio, R., Rode, G., & Bartolomeo, P. (2015). White matter lesional predictors of chronic visual neglect: a longitudinal study. Brain, 138(Pt 3), 746–760. doi: 10.1093/brain/awu389

Lupiáñez, J. (2010). Inhibition of return. In A. C. Nobre & J. T. Coull (Eds.), Attention and time (pp. 17–34): Oxford University Press.

Lupiáñez, J., Klein, R. M., & Bartolomeo, P. (2006). Inhibition of return: Twenty years after. Cognitive Neuropsychology, 23(7), 1003–1014.

Lupiáñez, J., Martín-Arévalo, E., & Chica, A. B. (2013). Is Inhibition of Return due to attentional disengagement or to a detection cost? The Detection Cost Theory of IOR. Psicológica, 34(2).

Lupiáñez, J., Milan, E. G., Tornay, F. J., Madrid, E., & Tudela, P. (1997). Does IOR occur in discrimination tasks? Yes, it does, but later. Perception and Psychophysics, 59(8), 1241–1254.

Macaluso, E., Frith, C. D., & Driver, J. (2000). Modulation of human visual cortex by crossmodal spatial attention. Science, 289(5482), 1206–1208.

Mars, R. B., Sallet, J., Schuffelgen, U., Jbabdi, S., Toni, I., & Rushworth, M. F. (2012). Connectivity-based subdivisions of the human right "temporoparietal junction area": evidence for different areas participating in different cortical networks. Cereb Cortex, 22(8), 1894–1903. doi: 10.1093/cercor/bhr268

Martín-Arévalo, E., Chica, A. B., & Lupiáñez, J. (2014). Electrophysiological modulations of exogenous attention by intervening events. Brain and cognition, 85, 239–250.

Mathôt, S., & Theeuwes, J. (2010). Gradual Remapping Results in Early Retinotopic and Late Spatiotopic Inhibition of Return. Psychological Science, 21(12), 1793–1798. doi: doi:10.1177/0956797610388813

Mort, D. J., Malhotra, P., Mannan, S. K., Rorden, C., Pambakian, A., Kennard, C., & Husain, M. (2003). The anatomy of visual neglect. Brain, 126(Pt 9), 1986–1997.

Nakamura, K., & Colby, C. L. (2002). Updating of the visual representation in monkey striate and extrastriate cortex during saccades. Proceedings of the National Academy of Sciences, 99(6), 4026–4031.

Patel, G. H., Yang, D., Jamerson, E. C., Snyder, L. H., Corbetta, M., & Ferrera, V. P. (2015). Functional evolution of new and expanded attention networks in humans. Proc Natl Acad Sci U S A, 112(30), 9454–9459. doi: 10.1073/pnas.1420395112

Pertzov, Y., Zohary, E., & Avidan, G. (2010). Rapid formation of spatiotopic representations as revealed by inhibition of return. Journal of Neuroscience, 30(26), 8882–8887.

Posner, M. I., & Cohen, Y. (1984). Components of visual orienting. In H. Bouma & D. Bouwhuis (Eds.), Attention and Performance X (pp. 531–556). London: Lawrence Erlbaum.

Posner, M. I., Rafal, R. D., Choate, L. S., & Vaughan, J. (1985). Inhibition of return: Neural basis and function. Cognitive Neuropsychology, 2, 211–228.

Robinson, D. L., Bowman, E. M., & Kertzman, C. (1995). Covert orienting of attention in macaques. II. Contributions of parietal cortex. Journal of Neurophysiology, 74(2), 698–712.

Robinson, D. L., & Kertzman, C. (1995). Covert orienting of attention in macaques. III. Contributions of the superior colliculus. Journal of Neurophysiology, 74(2), 713–721.

Röder, B., Spence, C., & Rösler, F. (2002). Assessing the effect of posture change on tactile inhibition-of-return. Experimental Brain Research, 143(4), 453–462.

Satel, J., Wang, Z., Hilchey, M. D., & Klein, R. M. (2012). Examining the dissociation of retinotopic and spatiotopic inhibition of return with event-related potentials. Neuroscience Letters, 524(1), 40–44.

Schettino, A., Rossi, V., Pourtois, G., & Müller, M. M. (2016). Involuntary attentional orienting in the absence of awareness speeds up early sensory processing. Cortex, 74, 107–117.

Schmahmann, J. D., & Pandya, D. N. (2006). Fiber Pathways of the Brain. New York: Oxford University Press.

Spence, C., Lloyd, D., McGlone F., Nicholls, M. E., & Driver, J. (2000). Inhibition of return is supramodal: a demonstration between all possible pairings of vision, touch, and audition. Experimental Brain Research, 134(1), 42–48.

Steinmetz, M., Connor, C., Constantinidis, C., & McLaughlin, J. (1994). Covert attention suppresses neuronal responses in area 7a of the posterior parietal cortex. Journal of Neurophysiology, 72(2), 1020–1023.

Swann, N. C., Tandon, N., Pieters, T. A., & Aron, A. R. (2012). Intracranial Electroencephalography Reveals Different Temporal Profiles for Dorsal- and Ventro-lateral Prefrontal Cortex in Preparing to Stop Action. Cerebral Cortex, 23(10), 2479–2488. doi: 10.1093/cercor/bhs245

Taylor, T. L., & Klein, R. M. (2000). Visual and motor effects in inhibition of return. Journal of Experimental Psychology: Human Perception and Performance, 26(5), 1639.

Thiebaut de Schotten, M., Dell’Acqua F., Forkel, S. J., Simmons, A., Vergani, F., Murphy, D. G. M., & Catani, M. (2011). A lateralized brain network for visuospatial attention. Nature Neuroscience, 14(10), 1245–1246. doi: 10.1038/nn.2905

Thiebaut de Schotten, M., Urbanski, M., Duffau, H., Volle, E., Levy, R., Dubois, B., & Bartolomeo, P. (2005). Direct evidence for a parietal-frontal pathway subserving spatial awareness in humans. Science, 309(5744), 2226–2228. doi: 10.1126/science.1116251

Thompson, K. G., & Bichot, N. P. (2005). A visual salience map in the primate frontal eye field. Progress in brain research, 147, 249–262.

Tipper, S. P., Rafal, R., Reuter-Lorenz, P. A., Starrveldt, Y., Ro, T., Egly, R., … Weaver, B. (1997). Object-based facilitation and inhibition from visual orienting in the human split-brain. Journal of Experimental Psychology: Human Perception and Performance, 23(5), 1522.

Tolias, A. S., Moore, T., Smirnakis, S. M., Tehovnik, E. J., Siapas, A. G., & Schiller, P. H. (2001). Eye movements modulate visual receptive fields of V4 neurons. Neuron, 29(3), 757–767.

Umeno, M. M., & Goldberg, M. E. (1997). Spatial processing in the monkey frontal eye field. I. Predictive visual responses. Journal of Neurophysiology, 78(3), 1373–1383.

Umeno, M. M., & Goldberg, M. E. (2001). Spatial processing in the monkey frontal eye field. II. Memory responses. Journal of Neurophysiology, 86(5), 2344–2352.

Vivas, A. B., Humphreys, G. W., & Fuentes, L. J. (2003). Inhibitory processing following damage to the parietal lobe. Neuropsychologia, 41(11), 1531–1540.

Vivas, A. B., Humphreys, G. W., & Fuentes, L. J. (2006). Abnormal inhibition of return: A review and new data on patients with parietal lobe damage. Cognitive Neuropsychology, 23(7), 1049–1064.

Wu, Y., Sun, D., Wang, Y., Wang, Y., & Wang, Y. (2016). Tracing short connections of the temporo-parieto-occipital region in the human brain using diffusion spectrum imaging and fiber dissection. Brain Research, 1646, 152–159. doi: http://dx.doi.org/10.1016/j.brainres.2016.05.046

Wurtz, R. H. (2008). Neuronal mechanisms of visual stability. Vision research, 48(20), 2070–2089.

Wurtz, R. H., Joiner, W. M., & Berman, R. A. (2011). Neuronal mechanisms for visual stability: progress and problems. Philosophical Transactions of the Royal Society of London B: Biological Sciences, 366(1564), 492–503.

Zipser, D., & Andersen, R. A. (1988). A back-propagation programmed network that simulates response properties of a subset of posterior parietal neurons. Nature, 331(6158), 679–684.

Zohary, E., Shadlen, M. N., & Newsome, W. T. (1994). Correlated neuronal discharge rate and its implications for psychophysical performance. Nature, 370(6485), 140–143.

